# Design of intrinsically disordered proteins that undergo phase transitions with lower critical solution temperatures

**DOI:** 10.1101/2020.11.13.381897

**Authors:** Xiangze Zeng, Chengwen Liu, Martin J. Fossat, Pengyu Ren, Ashutosh Chilkoti, Rohit V. Pappu

## Abstract

Many naturally occurring elastomers are intrinsically disordered proteins (IDPs) built up of repeating units and they can demonstrate two types of thermoresponsive phase behavior. Systems characterized by lower critical solution temperatures (LCST) undergo phase separation above the LCST whereas systems characterized by upper critical solution temperatures (UCST) undergo phase separation below the UCST. There is congruence between thermoresponsive coil-globule transitions and phase behavior whereby the theta temperatures above or below which the IDPs transition from coils to globules serve as useful proxies for the LCST / UCST values. This implies that one can design sequences with desired values for the theta temperature with either increasing or decreasing radii of gyration above the theta temperature. Here, we show that the Monte Carlo simulations performed in the so-called intrinsic solvation (IS) limit version of the temperature-dependent the ABSINTH (self-Assembly of Biomolecules Studied by an Implicit, Novel, Tunable Hamiltonian) implicit solvation model, yields a useful heuristic for discriminating between sequences with known LCST versus UCST phase behavior. Accordingly, we use this heuristic in a supervised approach, integrate it with a genetic algorithm, combine this with IS limit simulations, and demonstrate that novel sequences can be designed with LCST phase behavior. These calculations are aided by direct estimates of temperature dependent free energies of solvation for model compounds that are derived using the polarizable AMOEBA (atomic multipole optimized energetics for biomolecular applications) forcefield. To demonstrate the validity of our designs, we calculate coil-globule transition profiles using the full ABSINTH model and combine these with Gaussian Cluster Theory calculations to establish the LCST phase behavior of designed IDPs.

## Introduction

Intrinsically disordered proteins (IDPs) that undergo thermoresponsive phase transitions are the basis of many naturally occurring elastomeric materials ^1^. These naturally occurring scaffold IDPs ^2^ serve as the basis of ongoing efforts to design thermoresponsive materials ^3^. Well-known examples of disordered regions derived from elastomeric proteins ^4^, include the repetitive sequences from proteins such as resilins ^5^, elastins ^6^, proteins from spider silks ^7^, titin ^8^, and neurofilament sidearms ^9^. Elastin-like polypeptides have served as the benchmark systems for the development of responsive disordered proteins that can be adapted for use in various biotechnology settings ^10^. The multi-way interplay of sequence-encoded intermolecular interactions, chain-solvent interactions, as well as chain and solvent entropy gives rise to thermoresponsive phase transitions that lead to the formation of coacervates ^1^. Here, we show that one can expand the “materials genome” ^11^ through *de novo* design strategies that are based on heuristics anchored in the physics of thermoresponsive transitions and efficient simulation engines that apply the learned heuristics in a supervised approach. We report the development of a genetic algorithm (GA) and show how it can be applied in conjunction with multiscale computations to design thermoresponsive IDPs with LCST phase behavior.

Conformational heterogeneity is a defining hallmark of IDPs ^12^. Work over the past decade- and-a-half has shown that naturally occurring IDPs come in distinct sequence flavors ^12^. Indeed, IDPs can be distinguished based on their sequence-encoded interplay between intramolecular and chain-solvent interactions that can be altered through changes in solution conditions. Recent studies have shown that IDPs can be drivers or regulators of reversible phase transitions in simple and complex mixtures of protein and nucleic acid molecules ^13^. These transitions are driven primarily by the multivalence of interaction motifs that engage in reversible physical crosslinks ^14^. IDPs can serve as scaffolds for interaction motifs (stickers), interspersed by spacers. Alternatively, they can modulate multivalent interactions mediated by stickers that are situated on the surfaces ^15^ of autonomously foldable protein domains ^16^.

Thermoresponsive phase transitions arise either by increasing the solution temperature above a lower critical solution temperature (LCST) or by lowering the temperature below an upper critical solution temperature (UCST) ^1^. Many systems are capable of both types of thermoresponsive transitions, although only one of the transitions might be accessible in the temperature range of interest. Here, we leverage our working knowledge of the sequence features that encode driving forces for thermoresponsive phase transitions ^17^ to develop and deploy a GA for the design of novel IDPs characterized by LCST behavior. Inspired by work on elastin-like polypeptides ^3^, we focus on designing IDPs that are repeats of pentapeptide motifs. The amino acid composition of each motif contributes to the LCST behavior and the number of repeats determines the multivalence of stickers that drive phase transitions with LCST behavior.

The GA we adapt for this work is driven by advances that include: (a) an improved fundamental understanding of the physics of LCST phase behavior ^18^; (b) experiments showing that many IDPs undergo collapse transitions with increased temperature ^19^; (c) a generalization of the ABSINTH implicit solvation model and forcefield paradigm^20^ to account for the temperature dependence of chain solvation; (d) a growing corpus of information regarding the sequence determinants of LCST phase behavior in repetitive IDPs ^17^; and (e) the prior demonstration that a GA based method known as GADIS (Genetic Algorithm for Design of Intrinsic Secondary structure) ^21^ can be combined with efficient, ABSINTH-based simulations to design IDPs with bespoke secondary structural preferences.

Studies of synthetic polymer systems have helped in elucidating the origins of the driving forces for and the mechanisms of LCST phase behavior ^22^. A well-known example is poly-N-isopropylacrylamide (PNIPAM) ^23^. Here, the dispersed single phase is stabilized at temperatures below ∼32°C by the favorable hydration of amides in the sidechains. Solvation of amides requires that the solvent be organized around the hydrophobic moieties that include the backbone carbon chain and the isopropyl group in the sidechain. The entropic cost for organizing solvent molecules around individual chains increases with increasing temperature. Accordingly, above the LCST of ∼32°C, and for volume fractions that are greater than a threshold value, the system phase separates to form a polymer-rich coacervate phase that coexists with a polymer-poor dilute phase. The driving forces for phase separation are the gain in solvent entropy through the release of solvent molecules from the polymer and the gain of favorable inter-chain interactions, such as hydrogen-bonding interactions between amides in the polymer.

Tanaka and coworkers have developed a *cooperative hydration* approach, inspired by the Zimm and Bragg theories for helix-coil transitions ^24^, to model the physics of phase transitions with LCST ^25^. Cooperative hydration refers to the cooperative association (below the LCST) or dissociation (above the LCST) of water molecules that are bound to repeating units along the polymer chain ^26^. Cooperativity is captured using the Zimm-Bragg formalism by modeling each repeating unit as being in one of two states *viz*., solvated or desolvated. In the solvated state, the repeating unit has a defined interaction strength with solvent molecules. In the desolvated state, pairs of such repeating units have defined exchange interactions. In addition, desolvation is associated with a gain in solvent entropy. The three-way interplay of direct solvent-chain interactions, the interactions among desolvated pairs of units, and the gain in solvent entropy above the LCST can be captured in a suitable physical framework that can be parameterized to describe system-specific phase transitions. Accordingly, if one has prior knowledge of the interaction energies associated with each repeat unit, one can use the framework of Tanaka and coworkers to design novel sequences with LCST behavior.

An alternative approach, which we adopt in this work, is to leverage the corollary of LCST behavior at the single chain limit ^27^. At temperatures that are proximal to the LCST, the chain of interest will undergo a coil-to-globule transition in a dilute solution ^28^. This is because chain collapse is a manifestation of the physics of phase separation in the single chain limit. Here, we leverage this connection between phase separation and chain collapse of isolated polymer chains in ultra-dilute solutions to design novel IDPs that are predicted to undergo phase transitions with LCST phase behavior. We do so by using a multi-pronged approach that starts with improved estimates of the temperature dependencies of free energies of solvation of model compounds that mimic amino acid sidechain and backbone moieties. For this, we use free energy calculations based on the AMOEBA forcefield ^29^, which is a second-generation, molecular mechanics based, polarizable model for water molecules and proteins. We incorporate these temperature dependent free energies of solvation into the ABSINTH implicit solvation model and show that thermoresponsive changes to chain dimensions, calculated in the “*intrinsic solvation* (IS) limit” ^30^, yields robust heuristics that discriminates sequences with known LCST phase behavior from those that show UCST behavior. We then describe the development of a GA, an adaptation of the GADIS approach, to design novel sequences that relies on all-atom simulations, performed using the ABSINTH model in the IS limit, and learned heuristics as fitness scores. Distinct classes of designed sequences emerge from our approach and these are screened to filter out sequences with low disorder scores as assessed using the IUPRED2 algorithm ^31^. The resulting set of sequences are analyzed using simulations based on the full ABSINTH model, which show that the designed sequences do undergo collapse transitions above a threshold temperature. The contraction ratio, defined as the ratio of chain dimensions at temperature *T* to the dimensions at the theta temperature and computed as a function of simulation temperature, is analyzed to extract temperature dependent two-body interaction parameters and athermal three-body interaction parameters that are used in conjunction with the Gaussian Cluster Theory (GCT) ^32^ to calculate system-specific phase diagrams ^28^. The upshot is a multiscale pipeline whereby a GA, aided by a derived heuristic and IS limit simulations, leads to the design of novel sequences with predicted LCST phase behavior. Following a post-processing step that selects for sequences with a high confidence of being intrinsically disordered, we combine all-atom ABSINTH-T based simulations with Gaussian Cluster Theory to obtain sequence-specific phase diagrams. These last two steps allow further pruning of the sequence space derived from the designs and provide further confidence regarding the authenticity of the predicted LCST phase behavior.

Temperature-dependent free energies of solvation are central to accurate descriptions of LCST behavior. Each protein may be viewed as a chain of model compounds and measured / calculated temperature dependent values of temperature dependent free energies of hydration Δµ_h_ for fully solvated model compounds can be used as the reference free energies of solvation (rFoS) in implicit solvation models such as EEF1^33^ or ABSINTH ^20^. Where possible, the ABSINTH model ^20,34^ uses experimentally measured free energies of solvation for model compounds. In the original formalism, Vitalis and Pappu ^20^ adapted experimentally derived rFoS values at 298 K and assumed these values to be independent of temperature. This approach was generalized by Wuttke et al.,^19^ to calculate temperature dependent rFoS values, using data from calorimetric measurements made by Makhatadze and Privalov^35^ for the enthalpy and heat capacity of hydration at a reference temperature. These values were augmented by those of Cabani et al.,^36^ for naphthalene, which is used as a model compound mimic of tryptophan. Wuttke et al.,^19^ incorporated the enthalpy and heat capacity of hydration estimated at a reference temperature into an integrated version of the Gibbs-Helmholtz equation to yield a thermodynamic model for temperature dependent rFoS values for all the relevant model compounds. In this formalism, rFoS(*T*) or Δµ_h_(*T*) is written as:

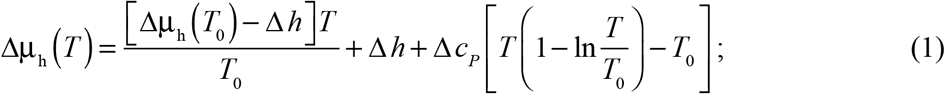

Here, Δ*h* is the enthalpy of solvation (hydration) at a reference temperature *T*_0_, which is typically set to be 298 K, and Δ*c*_*P*_ is the molar heat capacity change associated with the solvation process. Based on measurements, the assumption is that Δ*c*_*P*_ is independent of temperature ^37^.

We built on the approach of Wuttke et al.,^19^ to incorporate temperature dependent rFoS values in ABSINTH. This is implemented in a version that we refer to as ABSINTH-T. The issues we faced in developing ABSINTH-T were two-fold. First, the values for Δ*c*_*p*_ and Δ*h* that were used by Wuttke et al., rely on decompositions of measurements for model compounds into group-specific contributions. In contrast, the Δµ_*h*_values used in ABSINTH are for model compounds and explicitly avoid the group-specific decompositions made by Makhatadze and Privalov ^35^. This choice reflects the fact that group-specific decompositions are not measured. Instead, they are derived quantities that are based on empirical reasoning. This creates a mismatch with the paradigm that underlies the ABSINTH framework ^20^. Put simply, we require values of Δµ_h_, Δ*c*_*p*_ and Δ*h* that correspond to model compounds as opposed to group-specific decompositions.

Second, model compounds that mimic the sidechains of ionizable residues pose unique challenges. For any solute, including ions, the free energy of hydration at a specific temperature and pressure is defined as the change in free energy change associated with transferring the solute of interest from a dilute vapor phase into water ^38^. The accommodation of the solute into liquid water is associated with the cost to create a cavity in the solvent ^38^, the electronegative cavity potential ^39^, the work to add soft dispersion interactions ^40^, and distribute charges uniformly or non-uniformly across the solute ^41^. Vapor pressure osmometry with radioactive labeling, as used by Wolfenden^42^ to measure free energies of hydration for polar solutes, including neutral forms of ionizable species, cannot be used to measure free energies of hydration of ions because of the ultra-low vapor pressures and the confounding effects of ion-pairing in the gas phase. Calorimetry, as used by Makhatadze, Privalov, and colleagues provides an alternative approach ^35,43^. However, the large magnitudes of free energies of hydration, which are expected to be on the order 10^2^ kcal/mol ^44,45^, giving rise to even larger magnitudes for enthalpies of hydration, make it impossible to obtain the numbers of interest directly from calorimetric measurements. Measurements of activity coefficients on the concentration of whole salts in aqueous solutions can be used to place bounds on the values of Δµ_h_ ^46^, but these are not direct measurements of Δµ_h_.

A key challenge is that stable solutions are electroneutral ^45^. Accordingly, all measurements aimed at estimating the free energies of hydration of ionic species have to rely on parsing numbers derived from measurements on whole salts against those of reference salts ^44^– see the work of Grossfield et al.,^41^ and references therein. Alternatives rely on referencing measurements for whole salts against the free energy of hydration of the proton ^47^ – a quantity plagued by considerable uncertainty given the interplay between Zundel and Eigen forms for the hydronium ion ^48^. One can also use direct measurements of neutralized versions of ionic species ^35,42,43^; however, extracting the parameters of interest ends up relying on explicit or implicit assumptions regarding proton hydration free energies to extract estimates of the desired free energies of hydration of ionic species. The upshot is that direct measurements of free energies of hydration of ionic species are not feasible, and hence one has to rely on the validity of models that are used to parse experimental data.

In 1996, in their work aimed at accounting for reaction-field effects in calculations of hydration free energies in continuum models, Marten et al.,^49^ compiled a set of values for hydration free energies for all the relevant model compounds. In the original ABSINTH model, ^20^ the values tabulated by Marten et al., were used for all uncharged solutes. For charged species, specifically the protonated versions of Arg and Lys sidechains and deprotonated versions of Asp and Glu sidechains, Vitalis and Pappu ^20^ used the numbers tabulated and parsed by Marcus ^50^ for a reference temperature of 298 K.

Wuttke et al.,^19^ used the rFoS values tabulated by Vitalis and Pappu ^20^ at 298 K, and tested three different models for generating T-dependent rFoS values of model compounds used to mimic the charged versions of Arg, Asp, Lys and Glu. Model 1 uses the measured enthalpies and heat capacities measured for the neutral compounds ^35^, i.e., protonated Asp and Glu and deprotonated Lys and Arg. These were then scaled by the rFoS values used by Vitalis and Pappu ^20^ for the charged variants. The scaled enthalpies and heat capacities were then deployed in Equation (1). Model 2 of Wuttke et al.,^19^ uses the enthalpies of hydration estimated by Marcus ^51^ and the heat capacities of hydration tabulated by Abraham and Marcus ^52^. As noted above, these numbers are not direct measurements. Instead, they were derived from measurements of whole salts and then parsed using different models to arrive at a consensus set of estimates for the enthalpies and heat capacities. Model 3 of Wuttke et al.,^19^ uses the same heat capacities as model 2, and empirical choices were made for the enthalpies based on “expectations for a variety of charged model compounds”.

The preceding discussion emphasizes the fact that direct measurements of the rFoS values as a function of temperature or of the enthalpies and heat capacities of hydration at reference temperatures are unavailable for model compounds that mimic charged versions of the sidechains of Arg, Asp, Lys, and Glu. To put the challenge into perspective, we note that models 1 and 2 of Wuttke et al.,^19^ yield values of 50.37 cal mol^-1^K^-1^ and 5.30 cal mol^-1^K^-1^, respectively for the Δ*c*_*p*_ of the acetate ion. The large variations are a reflection of the challenges associated with estimating temperature independent and temperature rFoS values for charged species.

Here, we pursue a different approach: we use AMOEBA, which is a second generation molecular mechanics based polarizable forcefield ^29^, in direct calculations of T-dependent rFoS values for all the relevant model compounds. The AMOEBA water model reproduces the temperature-dependent anomalies of liquid water ^53^ and yields accurate free energies of solvation for model compounds in aqueous solvents ^29,54,55^. Our goal was to have a common source for T-dependent rFoS values of the key model compounds that are used in ABSINTH. The free energy calculations were performed at specific temperatures and the integrated version of the Gibbs-Helmholtz equation was used to the data to extract Δ*h* and Δ*c*_*P*_. The values of Δ*h* and Δ*c*_*P*_ in conjunction with Equation (1) are used to calculate T-dependent rFoS values in ABSINTH-T.

## Results and Discussion

### Results from AMOEBA-based free energy calculations for model compounds

We performed temperature dependent free energy calculations based on the Bennett Acceptance Ratio (BAR) free energy estimator ^56^ for direct investigation of how Δµ_h_ varies with temperature. These calculations were performed for nineteen different model compounds that mimic the twenty sidechain moieties and the backbone peptide unit. Details of the parameterization of the AMOEBA forcefield for model compounds used in this study, and the design of the free energy calculations are provided in the methods section.

The temperature-dependent values for Δµ_h_ with error bars are shown in **Table S1** of the *Supporting Information*. **Figure S1** shows two sets of plots that compare the AMOEBA-derived rFoS values at 298 K to direct measurements for uncharged molecules, and to inferred values from parsing of data for charged compounds. The calculated values are in good agreement with experimental data for uncharged molecules. This is reassuring because AMOEBA is parameterized directly from *ab initio* quantum mechanical calculations and no knowledge is used with regard to condensed phases or experimental data in condensed phases. We do observe deviations between the AMOEBA derived rFoS values of charged species and the inferred values from experimental data for whole salts (**Figure S1**). These deviations are in accord with the concerns expressed in the introduction. Inasmuch as the AMOEBA derived values are direct calculations, we use these numbers as a self-consistent set for uncharged and charged molecules alike.

Results from temperature dependent calculations of Δµ_h_ for the nineteen relevant model compounds are shown in **Figure 1**. The enthalpy of hydration (Δ*h*) at *T*_0_ = 298 K and the temperature independent heat capacities of hydration (Δ*c*_*P*_) were extracted for each model compound by fitting the calculated temperature dependent free energies of solvation to the integral of the Gibbs-Helmholtz equation. The results are summarized in **Table 1**. As expected ^37^, the large positive heat capacity of hydration combined with the favorable enthalpies and unfavorable entropies lead to non-monotonic temperature dependencies for model compound mimics of the sidechain moieties of Ala, Val, Leu, Ile, and Pro. Similar results are observed for mimics of Phe, Tyr, and Trp. Of import, are the differences in hydration thermodynamics of the model compounds that mimic sidechains of Lys, Arg, Asp, and Glu. The model compounds 1-butylamine and *n*-propylguanidine that mimic the sidechains of Lys and Arg feature a duality of favorable enthalpy of hydration and large positive values for Δ*c*_*P*_. Finally, the deprotonated versions of acetic acid and propionic acid that mimic the deprotonated versions of Asp and Glu, respectively, have the most favorable free energies of hydration across the temperature range studied. Interestingly, these two solutes stand out for their distinctive negative heat capacities of hydration. Inferences based on integral equation theories ^57^ suggest that negative heat capacities of hydration derive from a weakening of the favorable solute-solvent interactions and a reduction of the extent to which water molecules are orientationally distorted within and in the vicinity of the first hydration shell.

**Table 1:** Results from free energy calculations that summarize values obtained for Δµ_h_ at 298 K. Data for the temperature dependence of Δµ_h_ were fit to equation (1), setting T_0_ = 298 K, to extract values for Δ*h* and Δ*c*_*P*_.

| Residue / unit | Model compound | $\Delta\mu_h$<br>kcal/mol | $\Delta h$<br>kcal/mol | $\Delta c_P$<br>cal / mol-K |
| --- | --- | --- | --- | --- |
| Ala | methane | 1.63 | -2.57 | 48.93 |
| Val / Pro | propane | 1.85 | -6.33 | 105.80 |
| Leu | 2-methalpropane | 2.22 | -5.92 | 109.38 |
| Ile | *n*-butane | 2.00 | -6.34 | 105.23 |
| Met | ethyl methyl thioether | -1.92 | -10.11 | 71.10 |
| Phe | toluene | -0.17 | -8.68 | 102.24 |
| Cys | methanethiol | -1.04 | -5.84 | 43.61 |
| Tyr | p-Cresol | -5.85 | -15.62 | 71.09 |
| Trp | 3-Methylindole | -4.46 | -12.67 | 108.10 |
| Ser | methanol | -5.08 | -10.41 | 10.43 |
| Thr | ethanol | -4.98 | -12.55 | 50.06 |
| Asn | acetamide | -8.61 | -14.37 | 6.18 |
| Gln | propionamide | -8.39 | -16.06 | 51.47 |
| His | 4-methylimidazole | -10.04 | -17.60 | 38.01 |
| backbone / Gly | N-methylacetamide | -8.33 | -16.10 | 44.73 |
| Arg | *n*-propylguanidine | -47.62* | -57.24* | 69.39 |
| Lys | 1-butylamine | -60.49* | -70.37* | 29.98 |
| Asp | acetic acid | -89.91* | -98.65* | -44.97 |
| Glu | propionic acid | -86.16* | -96.62* | -8.75 |

\*As with the default ABSINTH model, in ABSINTH-T, the rFoS values, and therefore the $\Delta h$ values we used for ionizable residues are offset from the calculated $\Delta\mu_h$ by a fixed constant of -30 kcal/mol. This, as was shown in the original work, is required to avoid the chelation of solution ions around ionizable residues. This “feature” remains a continuing weakness of the ABSINTH paradigm and one that we hope to remedy through suitable generalization of the model used in ABSINTH to interpolate between fully solvated and fully desolvated states.

**Figure 1:**
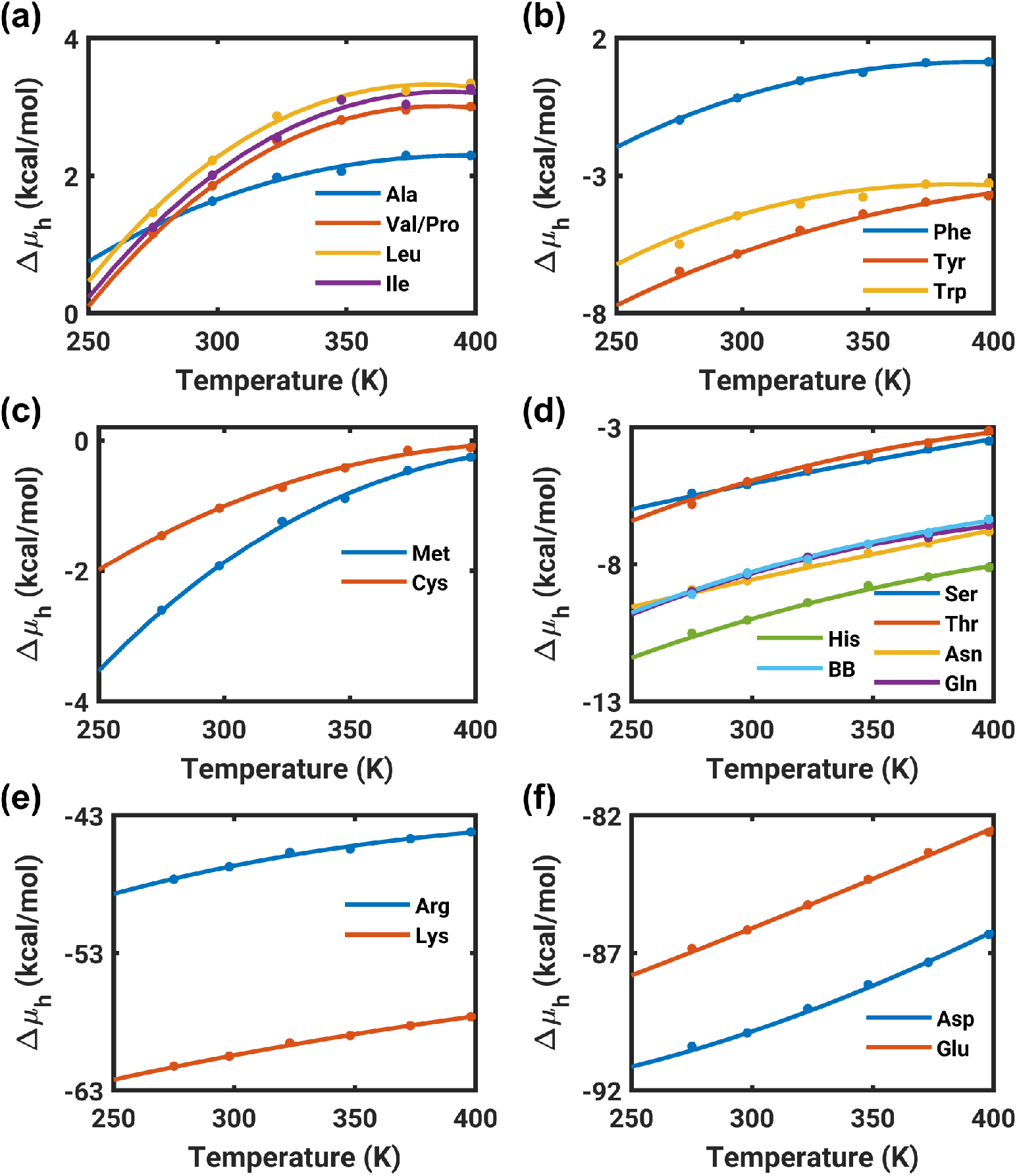
Temperature dependent free energies of solvation Δµ_h_ for model compounds that mimic sidechain and backbone moieties. The dots show results from free energy calculations based on the AMOEBA forcefield. These values are then fit to the integral of the Gibbs-Helmholtz equation (see main text) and the results of the fits are shown as solid curves. Parameters from the fits, which include estimates for Δ*h* and Δ*c*_*P*_ are shown in **Table 1**. In the legends we use the three letter abbreviations for each of the amino acids. Here, BB in panel (d) refers to the backbone moiety, modeled using N-methylacetamide, that mimics the peptide unit.

### Incorporation of T-dependent rFoS values into ABSINTH

In the ABSINTH model, each polyatomic solute is parsed into a set of solvation groups ^20,34^. These groups are model compounds for which the free energies of solvation rFoS are known *a priori*. In this work, we follow Wuttke et al.,^19^ and generalize the ABSINTH model to incorporate temperature dependencies of model compound rFoS values. In this ABSINTH-T model, the total solvent-mediated energy associated for a given configuration of the protein and solution ions is written as:

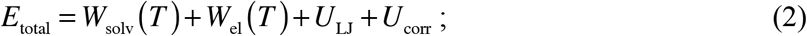

Here, *W*_solv_({rFoS(*T*)},{**r**}) is the many-body direct mean field interaction (DMFI) with the continuum solvent that depends on {rFoS(*T*)}, the set of temperature dependent rFoS values of model compounds that make up the solute and solution ions, and {**r**} is the set of configurational coordinates for polypeptide atoms and solution ions. The term *W*_solv_({rFoS(*T*)},{**r**})quantifies the free energy change associated with transferring the polyatomic solute into a mean field solvent while accounting for the temperature dependent modulation of the reference free energy of solvation for each solvation group due to other groups of the polyatomic solute as well as the solution ions. Additional modulations to the free energy of solvation of the solute due to interactions with charged sites on the polyatomic solute are accounted for by the *W*_el_ term. In ABSINTH-T the term *W*_el_({**r**},{**υ**(*T*)) is a function of the set of configurational coordinates {**r**}, solvation states {**υ**} of the solute atoms and solution ions, and the temperature dependent dielectric constant *ε*(*T*). For *ε*(*T*), we used the parameterization of Wuttke et al., ^19^. The effects of dielectric inhomogeneities, which are reflected in the configuration dependent solvation states, are accounted for without making explicit assumptions regarding the distance or spatial dependencies of dielectric saturation. The term *U*_LJ_ is a standard 12-6 Lennard-Jones potential and *U*_corr_ models specific torsion and bond angle-dependent stereoelectronic effects that are not captured by the *U*_LJ_ term. The ABSINTH paradigm is optimally interoperable with the OPLS-AA/L (Optimized Potentials for Liquid Simulations – All Atom / with LMP2 corrections) and the CHARMM ^58^ family of forcefields, and we use the OPLS-AA/L ^59^ forcefield.

### Intrinsic solvation (IS) approximation of ABSINTH-T as an efficient heuristic for discriminating IDPs with LCST versus UCST behavior

In the single chain limit, accessible in dilute solutions, polypeptides that show LCST phase behavior undergo collapse above a system specific theta temperature, whereas polypeptides that show UCST phase behavior expand above the system specific theta temperature ^1,28^. A GADIS-like strategy ^21^ for *de novo* design of polypeptide sequences with LCST phase behavior would involve ABSINTH-T based all-atom simulations to evaluate whether an increase in temperature leads to chain collapse. In effect, the fitness function in a GA comes from evaluation of the simulated ensembles as a function of temperature. Computationally, this becomes prohibitively expensive. Accordingly, we pursued a pared down version of ABSINTH-T, which is referred to as the intrinsic solvation (IS) limit of the model ^30^. The IS limit was introduced to set up sequence and composition specific reference models with respect to which one can use mean-field models to uncover how desolvation impacts IDP ensembles ^30,60^. In effect, the IS limit helps us map conformations in the maximally solvated ensemble and assess how this ensemble changes as a function of temperature. In the IS limit, the energy in a specific configuration for the sequence of interest is written as:

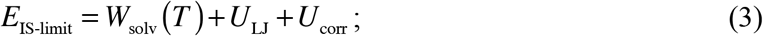

The only difference between the full model, see equation (2), and the IS limit is the omission of the *W*_el_ term. This increases the speed of simulations by 1-2 orders of magnitude depending on the system. Next, we asked if ensembles obtained from temperature dependent simulations performed in the IS limit could be used to obtain a suitable heuristic that discriminates sequences with LCST versus UCST behavior. These simulations were performed for a set of thirty sequences (see **Table S2** in the supporting information) that were previously shown by Garcia Quiroz and Chilkoti to have LCST and UCST phase behavior ^17^. The results are summarized in **Figure 2** and **Figures S2** and **S3** in the Supporting Information. As shown in panel (a) of **Figure 2**, the radii of gyration (*R*_*g*_), suitably normalized for comparisons across different sequences of different lengths, appear to be segregated into two distinct classes. To test this hypothesis, we computed the slopes *m* for each of the profiles of normalized *R*_*g*_ versus temperature. These slopes were calculated in the interval of simulation temperatures between 230 K and 380 K. The results, shown in panel (b) of **Figure 2**, clearly indicate that there indeed are two categories of sequences. Those that are known to show LCST phase behavior are colored in red, and they fall into a distinct group characterized by negative values of the slope *m* with an average value of –5.9 × 10^−3^ åK^−1^. Here, we use å to denote the units of *R*_*g*_ values normalized by the square root of the chain length *N*. In contrast, the slope for sequences that show UCST behavior is –1.4 × 10^−3^ åK^−1^. Given the range of sequences covered in the calibration based on the IS limit, we pursued an approach whereby we use slopes of *R*_*g*_ *N* ^−0.5^ versus *T* as a heuristic to guide the design of a genetic algorithm to find new sequences with LCST phase behavior. It is worth noting that we use the slopes of *R*_*g*_ *N* ^−0.5^ versus *T* plots instead of specific values of slopes of *R*_*g*_ *N* ^−0.5^ because: (a) *a priori* we would not know which temperature to choose for comparison of the *R*_g_ values; and (b) there is the formal possibility that the curves for *R*_*g*_ *N* ^−0.5^ versus *T* obtained for different constructs might cross one another, making the issue raised in (a) more confounding.

**Figure 2:**
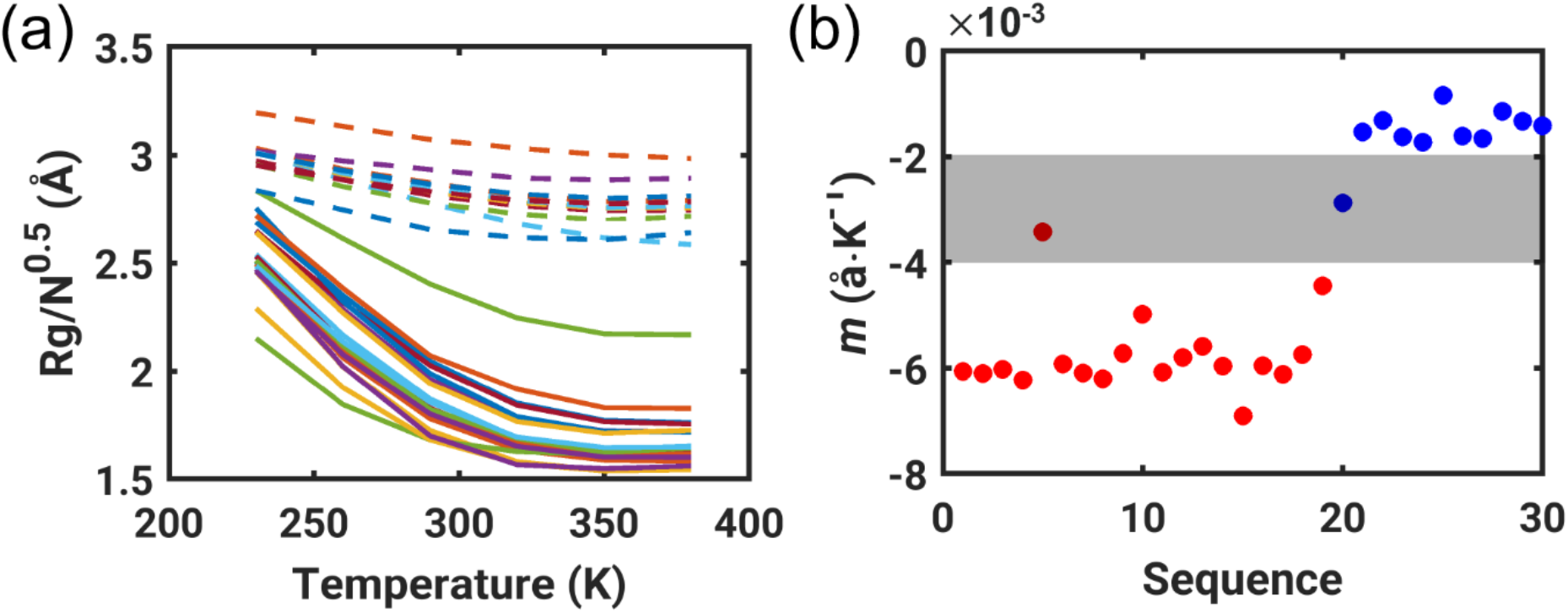
Analysis of IS limit simulations yields a heuristic that discriminates sequences with UCST vs. LCST phase behavior. (a) Plots of *R*_*g*_*N*^-0.5^ vs. temperature, extracted from IS limit simulations, for sequences shown by Garcia Quiroz and Chilkoti to have UCST (dashed lines) vs. LCST (solid lines) phase behavior. The sequences are shown in **Table S1** in the supporting information. (b) The slope *m* of the *R*_*g*_*N*^-0.5^ vs. temperature profiles. These slopes fall into two distinct categories, one for those with LCST phase behavior (blue) and another for those with UCST phase behavior (red). The gray region corresponds to the values of *m* that clearly demarcate the two categories of sequences.

### GA for the design of IDPs that are likely to have LCST phase behavior

We adapted the GADIS algorithm ^21^ to explore sequence space and discover candidate IDPs with predicted LCST phase behavior. To introduce the GA and demonstrate its usage, we set about designing novel sequences that are repeats of pentapeptide motifs. We focused on designing 55-mers, i.e., sequences with 11 pentapeptides. To keep the exercise simple, we focused on designing polymers that are perfect repeats of the pentapeptide in question. The GA used in this work is summarized in **Figure 3** and the details are described below.

**Figure 3.**
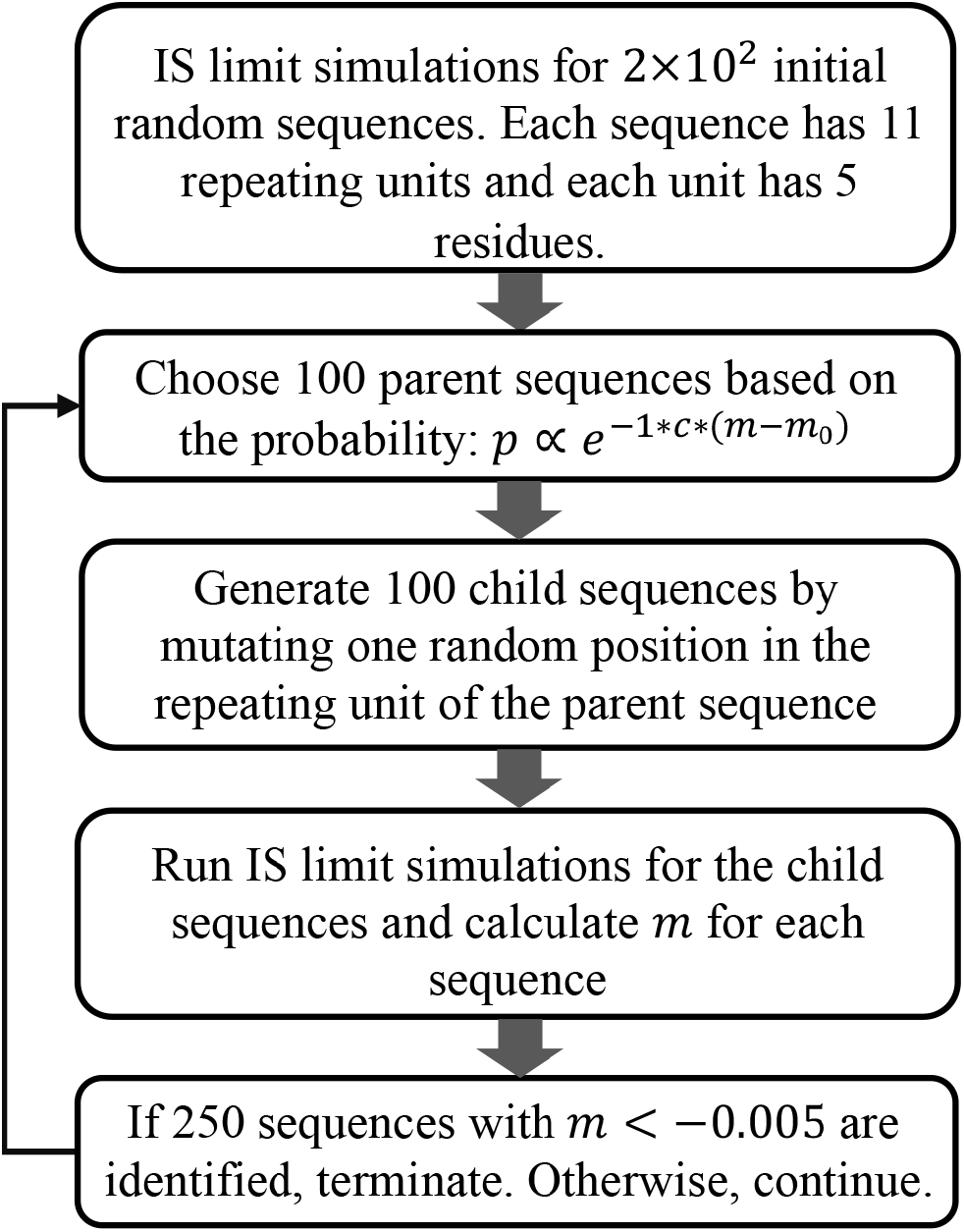
Workflow of the GA. We use this approach to design sequences that are predicted to have LCST phase behavior. A final post-processing step is added to filter our sequences that do not have high disorder scores (see main text).

The GA based design process is initiated by choosing a random set of 200 sequences. Next, for each of the random sequences we performed temperature based replica exchange ^61^ Metropolis Monte Carlo simulations in the IS limit. The simulation temperatures range from 200 K to 375 K with an interval of 25 K. From each converged IS limit simulation we computed the ensemble averaged *R*_*g*_ values as a function of simulation temperature *T*. These data were then used to evaluate the initial set of 200 values for the slope *m* using the following relationship:

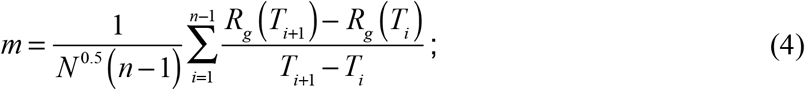

Here, *N* is the number of amino acids in each sequence, *n* is the number of replicas used in the simulation, and *T*_*i*_ is the temperature associated with replica *i*. The slope *m* was used to select 100 out of the 200 sequences that were chosen at random initially. The picking probability *p* was based on the following criterion:

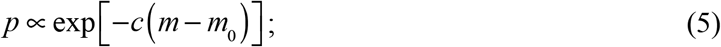

Here, *c* = 400 in units that are reciprocal to *m*, and *m*_0_ is set to –6.9 × 10^−3^ åK^−1^. This choice enables efficient evolution of the GA and a strong selection for sequences with negative values of *m*. The parameter *c* ensures numerical stability, guarding against the unnormalized value of *p* becoming too large or too small.

The chosen parent sequences were used to generate 100 child sequences by mutating a single, randomly chosen position to a randomly chosen residue in the repeating unit. To avoid the prospect of introducing spurious disulfide bonds, we do not include Cys residues either in the original parent pool or for propagating the child sequences. The GA was allowed to evolve for multiple iterations until the convergence criteria were met. These include the generation of at least 250 new sequences, each with a value of *m* being less than –5.0 × 10^−3^ åK^−1^. For the results presented here, six iterations were sufficient to meet the prescribed convergence criteria. The picking probability *p* determines the selection pressure encoded into the GA. There needs to be an optimal balance between the two extremes in selection pressure. High selection pressures can lead to early convergence to a local optimum whereas low selection pressures can drastically slow down convergence ^62^. The use of a single evolutionary operator can lead to a single sequence becoming the dominant choice. The number of iterations that pass before the emergence of a single sequence is known as the takeover time ^62^. High selection pressures lead to low takeover times and vice versa. The issue of a single dominant individual emerging is less of a concern in sequence design given the high dimensionality of sequence space. We tuned the choices for *c* and *m*_0_ to ensure that candidate sequences with putative UCST phase behavior can be part of the offspring, thus lending diversity to sequence evolution by the GA.

Panel (a) in **Figure 4** quantifies the progress of the GA through each iteration of the design process. The quantification is performed in terms of cumulative distribution functions, which for each iteration will quantify the probability that the emerging sequences have associated slope values that are less than or equal to a specific value. The rightward shift in each iteration is indicative of the improved fitness vis-à-vis the selection criterion, which is the lowering of *m*.

**Figure 4.**
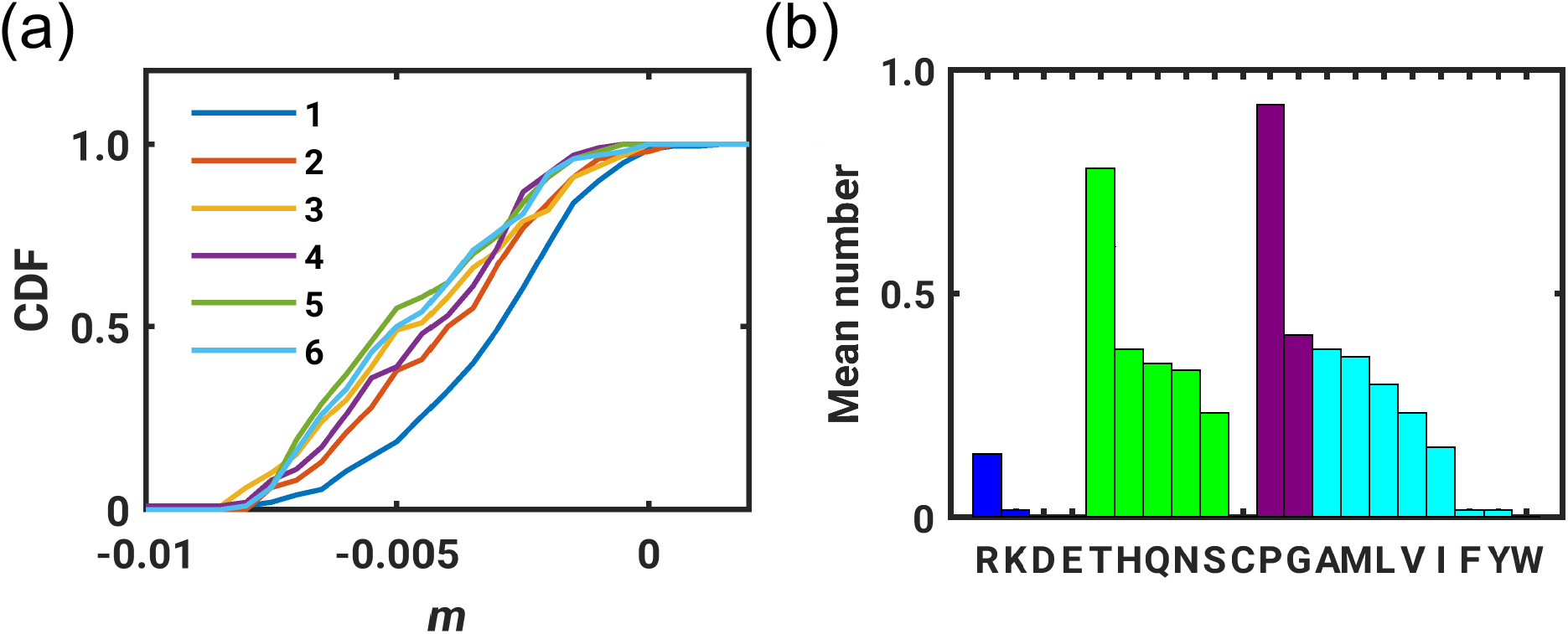
Calibration of the performance of the GA and statistics for compositional biases that emerge from application of the design protocol. (a) The cumulative distribution function (CDF) of the slope for sequences in each iteration. There is an overall shift for these CDFs towards smaller *m*-values with each iteration of the GA. (b) The mean number of each residue in the 64 designed IDPs that are predicted to show LCST phase behavior. Residues in panel (b) are grouped into categories based on their sidechain chemistries i.e., basic residues in blue bars, acidic residues in red bars (although these are not visible since they are not selected), polar residues in green, Pro and Gly in purple, and aliphatic as well as aromatic residues in cyan. Within each group, the bars are sorted in descending order of the mean numbers of occurrences in the designs.

Finally, we added a post-processing step to increase the likelihood that the designed sequences are *bona fide* IDPs. We used the disorder predictor IUPRED2 ^31^ to quantify disorder scores for each of the designed sequences. IUPRED2 yields a score between 0 and 1 for each residue, and only sequences where over half of the residues in the repeat are above 0.5 were selected as the final set of designed IDPs that are predicted to have LCST phase behavior. A particular concern with designing sequences for experimental prototyping is the issue of aggregation / precipitation. To ensure that designs were unlikely to create such problems, we calculated predicted solubility scores using the CamSol program ^63^ and found that all sequences that were selected after the post-processing step also have high solubility scores. This provides confidence that the designed IDPs are likely to show phase behavior via liquid-liquid phase separation above system-specific LCST values without creating problems of precipitation / aggregation.

Panel (b) in **Figure 4** summarizes the mean number of each amino acid type observed across the final tally of 64 designed sequences that survive the post-processing step. These statistics are largely in accord with the observations of Garcia Quiroz and Chilkoti ^17^. Essentially every sequence has at least once Pro residue in the repeat. The beta branched polar amino acid Thr is the other prominent feature that emerges from the selection. The remaining selection preferences fall into four distinct categories that include: (i) a clear preference for at least one polar amino acid *viz*., His, Ser, Thr, Asn, and Gln; (ii) a clear preference for the inclusion of at least one hydrophobic amino acid *viz*., Ala, Ile, Met, and Val; (iii) negligible selection, essentially an avoidance of the acidic residues Asp and Glu, as well as the aromatic residues Phe, Trp, and Tyr; and finally (iv) a weak preference for Arg over Lys, which is concordant with the distinct temperature dependent profiles for Δµ_h_ (**Figure 1**) and the large positive heat capacity of Arg (**Table 1**). Interestingly, if we fix the positions of Pro and Gly and select for residues in XPXXG or other types of motifs that are inspired by previous work on elastin-like polypeptides, the design process often converges on repeats that are known to be generators of polypeptides with *bona fide* LCST phase behavior (see **Figure S4** in the *Supporting Information*). This observation, and the statistics summarized in **Figure 4b** indicate that the design process uncovers sequences that are likely to have LCST phase behavior.

#### The designed sequences fall into distinct sequence classes

To quantify the degree of similarity among the set of designed sequences, we computed pairwise Hamming distances between all pairs of the 64 sequences. The resulting Hamming distances were then sorted, and sequences were clustered into distinct groups. Highly similar sequences have low Hamming distances, whereas the converse is true for dissimilar sequences. The resultant Hamming distance map is shown in **Figure 5**. The 64 sequences are unevenly distributed across nine major clusters. The actual sequences of the repeats, color-coded by their Hamming distance-based groupings, are shown in **Figure 6**. There are two features that stand out. First, sequences deviate from being repeats of VPGVG, which is the elastin-like motif. Second, we find that different sequence permutations on identical or similar composition manifolds emerge as candidates for LCST phase behavior. This observation suggests that at least in the IS limit it is the composition of each motif rather than the precise sequence that underlies adherence to the selection pressure in the GA. Interestingly, our observations are in accord with results from large-scale in vitro characterizations of sequences with LCST phase behavior ^64^. These experiments show that composition, rather than the precise sequence, is a defining feature of LCST phase behavior – a feature that is distinct from sequences that show UCST phase behavior ^3^.

**Figure 5.**
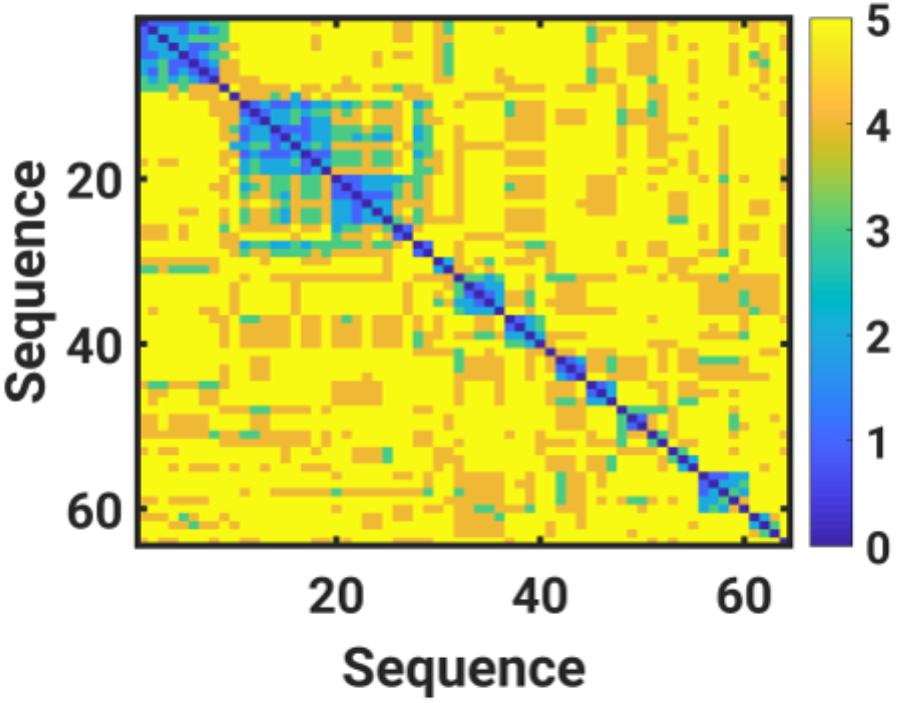
Identification of distinct sequence classes using a Hamming distance-based assessment of pairwise sequence similarities.

**Figure 6.**
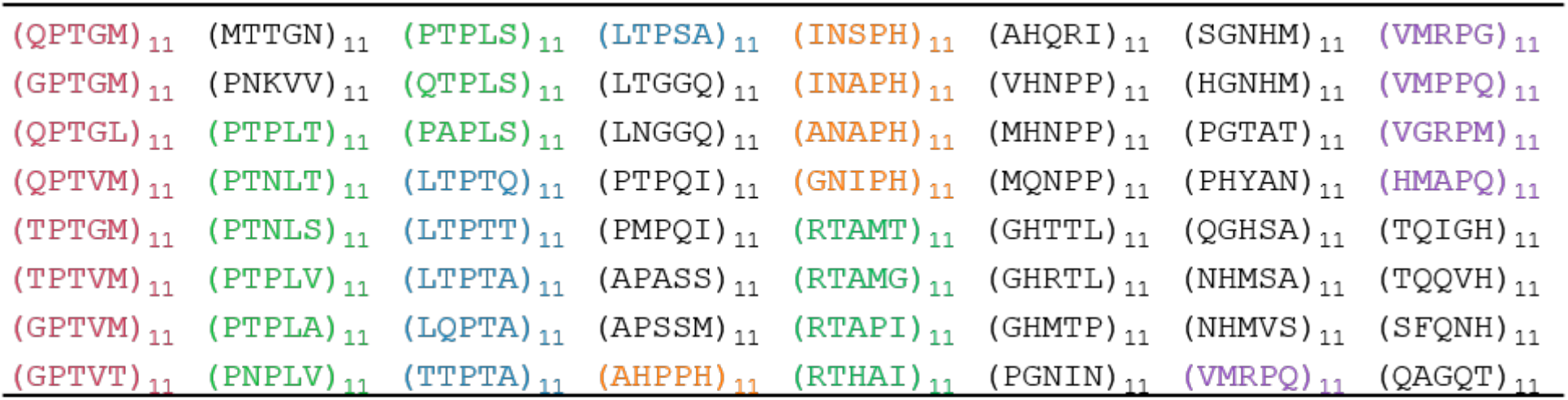
Sequences of 64 designed IDPs that emerge from application of the GA. Different colors except black are used to label sequences in the same group.

#### ABSINTH-T simulations of coil-to-globule transitions for select sequences

We selected four sequence repeats *viz*., (TPTGM)_11_, (PTPLV)_11_, (LTPTA)_11_, and (RTAMG)_11_ for characterization using the full ABSINTH-T model and the calculation of phase diagrams. These sequences were chosen because they are representatives from each of the four major classes that emerge from the design process. Additionally, these sequences bear minimal resemblance to extant designs or naturally occurring sequences that are known to have LCST phase behavior.

Using all-atom, thermal replica exchange Monte Carlo simulations and the full ABSINTH-T model we performed simulations to test for the presence of a collapse transition for each of the four sequences. The results are shown in **Figure 7**. All sequences show a clear tendency to form collapsed conformations as temperature increases. This is diagnosed by there being a clear preference for values of *R*_*g*_*N*^*-*0.5^ being less than the theta state reference value of 2.5 at higher temperatures and values of *R*_*g*_*N*^-0.5^ being greater than 2.5 at lower temperatures.

**Figure 7.**
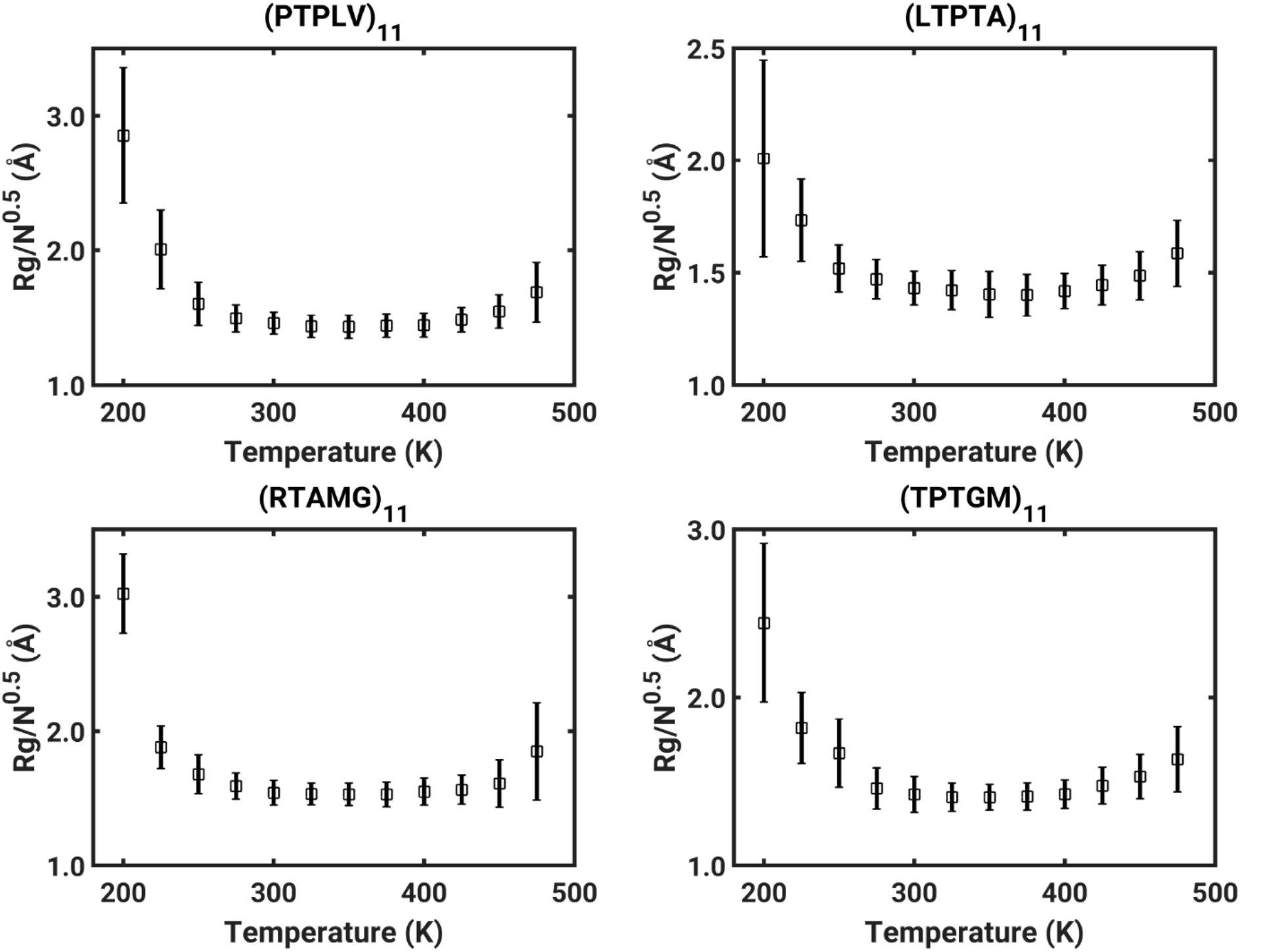
Profiles of normalized *R*_*g*_*N*^-0.5^ vs. temperature for four IDPs designed using the GA. The results shown here use the full ABSINTH-T model. The theta temperatures extracted from these simulations are presented in the main text.

#### Analysis of coil-globule transitions, extraction of parameters, and calculation of phase diagrams using the Gaussian Cluster Theory

The profiles of *R*_*g*_*N*^-0.5^ versus *T* were analyzed to extract the theta temperature (*T*_θ_) for each of the four sequences. For this, we used a method that described recently by Zeng et al., ^28^. Only three of the four sequences have coil-globule transition profiles for which a robust estimate of the theta temperature can be made. The extent of expansion at low temperatures is modest and suggests that the apparent *T*_θ_ for (LTPTA)_11_ is outside the window where converged simulations can be performed. For the other three sequences namely, (PTPLV)_11_, (RTAMG)_11_, and (TPTGM)_11_, the estimated *T*_θ_ values are 210 K, 210 K, and 200 K, respectively.

Next, we used the estimates of *T*_θ_ in conjunction with the Gaussian Cluster Theory of Raos and Allegra ^32^. We extracted the two and three-body interaction coefficients by fitting the contraction ratio *α*_s_ calculated from simulations using the formalism of the Gaussian Cluster Theory and this yields sequence-specific estimates of *B*, the two-body interaction coefficient, and *w*, the three-body interaction coefficient (see panels (a) – (c) in **Figure 8**). These parameters were then deployed to compute full phase diagrams using the numerical approach developed by Zeng et al.,^28^ and adapted by others ^65^. The results are shown in panels (d) – (f) of **Figure 8**. The abscissae in these diagrams denote the bulk polymer volume fractions whereas the ordinates quantify temperature in terms of the thermal interaction parameter 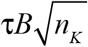 Here, 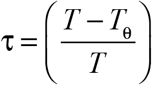 which is positive for *T* > *T*_θ_, *B* is the temperature-dependent two-body interaction coefficient inferred from analysis of the contraction ratio, and *n*_*K*_ is the number of Kuhn segment in the single chain, which we set to 8. Note that *B* is negative for temperatures above *T*_θ_. Accordingly, the thermal interaction parameter is positive above *T*_θ_ as well as the critical temperature *T*_c_. Therefore, comparative assessments of the driving forces for LCST phase behavior can be gleaned by comparing the sequence-specific values of 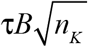 and the volume fraction at the critical point. It follows that the sequences can be arranged in descending order of the driving forces as (TPTGM)_11_, (RTAMG)_11_, and (PTPLV)_11_, respectively. Importantly, full characterization of the phase behavior using a combination of all-atom simulations and numerical adaptation of the Gaussian Cluster Theory shows that, in general, sequences designed to have LCST phase behavior, do match the predictions (see **Figure 8**).

**Figure 8.**
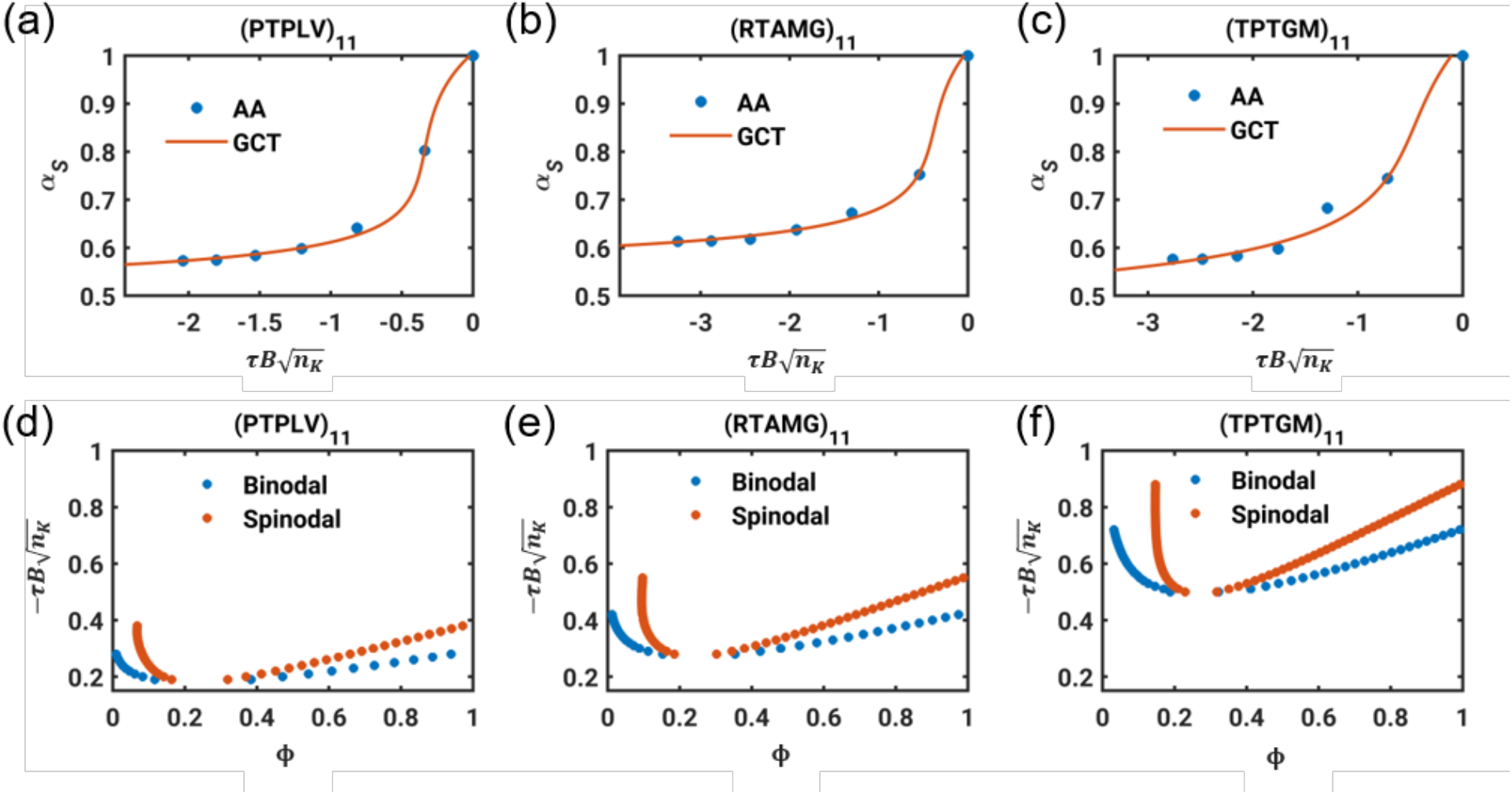
Results from application of the Gaussian Cluster Theory for calculating full phase diagrams. Panels (a-c) show the contraction ratio profiles for (PTPLV)_11_, (RTAMG)_11_ and (TPTGM)_11_, respectively. Blue dots are the contraction ratios calculated from all atom simulations with ABSINTH-T at temperatures from 200 K to 350 K and red curves are fits to these data using the Gaussian Cluster Theory that lead to estimates of the sequence-specific values for the temperature dependent two-body interaction coefficient *B* and the temperature independent three-body interaction parameter *w*. Panels (d-f) show the full phase diagrams, including the binodal, spinodal, and the estimated location of the critical point for (PTPLV)_11_, (RTAMG)_11_ and (TPTGM)_11_, respectively.

## Discussion

In this work, we have adapted a GA to design novel sequences of repetitive IDPs that we predict to have LCST phase behavior. Our method is aided by a learned heuristic that was shown to provide clear segregation between sequences with known LCST vs. UCST phase behavior. This heuristic is the slope *m* of the change in *R*_*g*_*N*^-0.5^ versus *T* from simulations of sequences performed in the IS limit of the ABSINTH-T model. We use the heuristic in conjunction with IS limit simulations to incorporate a selection pressure into the GA, thereby allowing the selection of sequences that are “fit” as assessed by the heuristic to be predictive of LCST phase behavior.

Here, we presented one instantiation of the GA and used it to uncover 64 novel sequences that can be grouped into four major classes and several minor classes (**Figure 6**). We then focused on four sequences, one each from each of the four major classes and characterized temperature dependent coil-globule transitions. These profiles, analyzed in conjunction with recent adaptations of the Gaussian Cluster Theory ^32^, allowed us to extract sequence-specific values for theta temperatures, temperature dependent values of the two body interaction coefficients, and three-body interaction coefficients. We incorporated these parameters into our numerical implementation ^28^ of the Gaussian Cluster Theory to calculate full phase diagrams for three sequences. These affirm the predictions of LCST phase behavior and demonstrates sequence-specificity in control over the driving forces for thermoresponsive phase behavior.

Our overall approach is aided by the following advances: We used the AMOEBA forcefield ^29^ to obtain direct estimates of temperature dependent free energies of solvation for model compounds used to mimic sidechain and backbone moieties. These temperature dependent free energies of solvation were used in conjunction with the integral of the Gibbs-Helmholtz equation to obtain model compound specific values for the enthalpy and heat capacity of hydration.

The methods we present here are a start toward the integration of supervised learning to leverage information gleaned from systematic characterizations of IDP phase behavior and physical chemistry based computations that combined all-atom simulations with improvements such as ABSINTH-T, and theoretical calculations that allow us to connect single chain coil-globule transitions to full phase diagrams ^28^. The heuristic we have extracted from IS limit simulations helps with discriminating sequences with LCST versus UCST phase behavior. These simulations are sufficient for IS limit driven and GA aided designs of sequences that are expected to have LCST phase behavior. This is because composition as opposed to the syntactic details of sequences play a determining role of LCST phase behavior ^3^. Recent studies have shown that even the simplest changes to sequence syntax can have profound impacts on UCST phase behavior ^66^. This makes it challenging to guide the design of sequences with predicted UCST phase behavior that relies exclusively on IS limit simulations. We will need to incorporate simulations based on either transferrable ^67^ or learned coarse-grained models ^68^ as a substitution for the IS limit simulations. This approach comes with challenges because one has to be sure that the coarse-grained models afford the requisite sequence specificity without compromising efficiency. The work of Dignon et al., ^69^ is noteworthy in this regard. Their coarse-grained model, which is based on knowledge-based potentials parameterized to have temperature-dependent interactions, have been shown to be very effective in discriminating sequences that are shown to have UCST versus LCST phase behavior ^69^. The conceptual underpinnings of their approach and that presented here derive from the work of Wuttke et al.,^19^. It would be interesting to combine or compare our approach to that of Dignon et al., in the context of designing novel IDPs and characterizing their phase behavior. We view these approaches as being complementary rather than competing ones and we expect that the approaches will have distinct advantages in different settings. The specific feature of our approach is that the calculations, at least for designing sequences with LCST phase behavior, do not ever become more complex than single chain simulations. This has value for achieving design objectives. It also has value for designing sequences that are not only thermoresponsive, but are also responsive to changes in pH, pressure, and other solvent parameters, especially since recent studies suggest that solution space scanning is a way to obtain efficient delineation of the desirable conformational and phase equilibria for IDPs ^70^.

The design of sequences with UCST phase behavior or sequences that combine UCST and LCST phase behavior, going beyond simple block copolymeric designs, will be of utmost interest for developing new IDP based materials. Additionally, we hope to build on improved understanding ^71^ of the impact of pH on conformational ^72^ and phase equilibria ^73^ of IDPs as well as the impact of metal chelation sites on phase behavior ^74^ to design sequences that combine the ability to exhibit phase behavior in response to orthogonal stimuli. Such efforts are of direct relevance to engineering orthogonal biomolecular condensates into simple unicellular prokaryotic and eukaryotic cells, as has been demonstrated recently with the engineering a protein translation circuit into protocells based on a thermoresponsive elastin like polypeptide ^75^. Of course, the proof of the validity / accuracy of designs and predictions will have to come from experimental work geared toward testing the predictions / designs. These efforts – that leverage high-throughput expression of these *de novo* sequences in *E. coli* and *in situ* characterization of their phase behavior – are underway ^76^. Initial experimental investigations suggest that the designs reported here and those that will emerge from application of the methods deployed in this work do indeed show LCST phase behavior. Detailed reports of these experimental characterizations will follow in separate work.

## Methods

### AMOEBA force field parametrization for the model compounds of interest

To obtain values of free energies of solvation from AMOEBA simulations, we first derived the AMOEBA force field parameters for the model compounds listed in **Table 1** of the main text. The parameters for N-methylacetamide, methane, methanol, ethanol, toluene and p-Cresol are taken from previous work ^55^, which was released in the amoeba09.prm parameter file in the TINKER package ^77^. The parameters for other model compounds are derived following the standard automated protocol that has been established for the AMOEBA forcefield ^78^. Briefly, the protocol involves the following steps: Quantum chemical calculations were utilized to derive the electrostatic parameters; these include atom-centered partial charges, dipole and quadrupole moments. The molecular structures were fully optimized at MP2/6-31G* level of theory ^79^ followed by MP2/cc-pvtz calculations to obtain the electron density of the molecules. Then initial multipole parameters were determined via distributed multipole analysis calculation via GDMA (Gaussian Distributed Multipole Analysis) program ^80^. With the charges being fixed, the dipole and quadrupole moments were further fit to the electrostatic potential generated at MP2/aug-cc-pvtz level on a grid of points outside of the molecules, where the least square restrained optimization was used to keep the multipole moments close to their DMA (Distributed Multipole Analysis) derived values while providing improved electrostatic potentials. The `poledit` and `potential` programs of TINKER package ^77^ were used in this process.

The Thole damping ^81^ value of 0.39 and the standard AMOEBA atomic polarizabilities were assigned for each atom. Valence and van der Waals (vdW) parameters were directly assigned from the existing small molecule library and MM3 (Molecular Mechanics 3) force field, and the equilibrium values for bond lengths and bond angles were calculated from above QM-optimized geometry. Torsional parameters of rotatable bonds were obtained by comparing the conformational energy profile of QM and AMOEBA model, which includes electrostatics, polarization, vdW and valence terms. The dihedral angle was scanned by minimizing all torsions about the rotatable bond of interest at 30° intervals with restrained optimization at HF/6-31G* level of theory. The QM conformational energy was obtained as the single point energy at ωB97XD/6-311++G(d,p) level of theory ^82^. Torsions about the same rotatable bond that are also in-phase are collapsed into one set of parameters for the fitting, and the contributions are distributed evenly among the parameters. AMOEBA uses the traditional Fourier expansion up to six-fold. Here, the force constant parameters were fit using 1, 2 and 3-fold trigonometric forms. All the quantum calculations were performed using the Gaussian 09 software package ^83^. The parametrization process has been automated in the Poltype (version 2) software ^78^. All the parameters derived above are appended as part of a separate text file in the supporting information.

### Set up of molecular dynamics simulations using AMOEBA

All AMOEBA simulations were performed using the TINKER-OpenMM package ^84^. Each model compound was solvated in a cubic water box with periodic boundary conditions. The initial dimensions of the central cell were set to be 30×30×30 Å^3^. Following energy minimization, molecular dynamics simulations were performed using reversible reference system propagator algorithm integrator ^85^ with an inner time step of 0.25 ps and an outer time step of 2.0 fs in isothermal-isobaric ensemble (NPT) ensemble with the target temperature being between 273 and 400 K depending on the temperature of interest and the target pressure being 1 bar. The temperature and pressure were controlled using a stochastic velocity rescaling thermostat ^86^ and a Monte Carlo constant pressure algorithm ^87^, respectively. The particle mesh Ewald (PME) method^88^ with PME-GRID being 36 grid units, an order 8 *B*-spline interpolation ^89^, with a real space cutoff of 7 Å was used to compute long-range corrections to electrostatic interactions. The cutoff for van der Waals interactions was set to be 12 Å. This combination of a shorter cutoff for PME real space and longer cutoff for Buffered-14-7 potential has been verified ^90^ for AMOEBA free energy simulations ^91^. Snapshots were saved every ps. In simulations performed along a prescribed schedule for the Kirkwood coupling parameters (please see below), we use the same solvent box across the schedule. However, the velocities were randomized at the start of each simulation, and the first 1 ns of data were set aside as equilibration, and not used in the free energy estimations.

### Free energy calculations

We used the Bennett Acceptance Ratio (BAR)^56^ method to quantify the free energies of solvation for the model compounds of interest. This method has been shown to be superior to other free energy estimators in terms of reducing the statistical errors in calculations of free energies of solvation ^92^. The solute is grown in using two different Kirkwood coupling parameters λ_vdW_ and λ_el_ that scale the strengths of solute-solute and solute-solvent van der Waals and electrostatic interactions. A series of independent molecular dynamics simulations were performed in the NPT ensemble for different combinations of λ_vdW_ and λ_el_. A soft-core modification of the Buffered-14-7 function was used to scale the vdW interactions as implemented in Tinker-OpenMM ^84^. We used the following combinations for the scaling coefficients: [λ_vdW_, λ_el_] λ ≡ [0, 0], [0.1, 0], [0.2, 0], [0.3, 0], [0.4, 0], [0.5, 0], [0.6, 0], [0.7, 0], [0.8, 0], [0.9, 0], [1, 0], [1, 0.1], [1, 0.2], [1, 0.3], [1, 0.4], [1, 0.5], [1, 0.6], [1, 0.7], [1, 0.8], [1, 0.9], [1, 1]. For each pair of λ values, we performed simulations, each of length 6 ns, at the desired temperature and a pressure of 1 bar. We then used the TINKER `bar` program to calculate the free energy difference between neighboring windows defined in terms of the scaling coefficients. For every combination of λ_vdW_ and λ_el_, we set aside the first 1 ns simulation as part of the equilibration process. Finally, for each model compound we computed free energies of solvation at six different temperatures *viz*., 275 K, 298 K, 323 K, 348 K, 373 K, 398 K, thus giving us the direct estimates of temperature dependent free energies of solvation that we sought from the AMOEBA based simulations. Note that 398 K is above the boiling point of water. However, although the physical properties of water are accurately captured by the AMOEBA model, the finite size of the system, the starting conditions, and the finite duration of the simulations, even though they are in the NPT ensemble, imply that water at 398 K and 1 bar corresponds to superheated liquid water.

The temperature dependent free energies of solvation were fit to the integral of the Gibbs-Helmholtz equation – see equation (1) in the main text. The free energy calculations provide us with direct estimates for rFoS(*T*) at specific values for *T*. We set *T*_0_ = 298K and fit use non-linear regression to fit equation (1) to the calculated values for rFoS(*T*). The regression analysis provides estimates of Δ*h* and Δ*c*_*P*_, which we then use, in conjunction with equation (1) in the manner prescribed by Wuttke et al.,^19^ for all the ABSINTH-T based simulations.

### Setup of Monte Carlo simulations in the IS limit and using ABSINTH-T

Thermal replica exchange ^61^ Monte Carlo simulations were performed using version 2.0 of the CAMPARI modeling software (http://campari.sourceforge.net/). The temperature schedule for thermal replica exchange simulations that use the full ABSINTH-T model ranges from 200 K to 470 K with an interval of 25 K. A total 6×10^7^ independent moves were attempted per replica. For systems in **Figure 7**, we performed three independent sets of thermal replica exchange simulations. All the simulations are performed within a spherical droplet with the radius of 100 Å. The other settings were identical to those used by Zeng et al., ^28^.

Details of the simulations including parameters, move sets, analyses, and design of the simulations are identical to those published in the recent work of Zeng et al., ^28^. Briefly, we used the ABSINTH-T implicit solvent model and forcefield paradigm. The forcefield parameters are based on the `abs_opls_3.2.prm` set and they include the parameters for proline residues that were developed by Radhakrishnan et al., ^93^. However, they do not include the CMAP corrections introduced by Choi and Pappu ^34^. The AMOEBA-based rFoS values at 298 K were incorporated into the standard parameter file, and the temperature dependent rFoS values were calculated using the model compound specific values for Δ*h* and Δ*c*_*p*_ that were derived using the AMOEBA-based calculations of Δµ_h_(*T*_0_) – see **Table 1**. As in our recent work ^28^, we used the temperature dependent dielectric constant prescribed by Wuttke et al., ^19^. Neutralizing counterions were added to the simulation droplet for polypeptides with net charges to neutralize the system. For the Na^+^ and Cl^-^ ions we use the following values for Δµ_h_ at 298 K, Δ*h*, and Δ*c*_*p*_, respectively: {-74.6 kcal / mol, -80.2 kcal / mol, -18.4 cal/mol-K} and {-87.2 kcal / mol, -99.2 kcal / mol, -11.7 cal/mol-K}. The Lennard-Jones parameters for Na^+^ and Cl^-^ ions are default parameters in the original work of Vitalis and Pappu ^20^.

For the IS limit simulations, we turned off the *W*_el_ term by setting the keyword SC_POLAR to be 0 in the key file. For each of the systems shown in **Figure 2**, we performed one set of replica exchange simulations, and the total of 6 × 10^7^ independent moves were attempted per replica. The temperature schedule for the replica exchange simulation is from 230 K to 380 K with an interval of 30 K. Error bars in **Figure 7** as well as **Figures S2** and **S3** are reported as standard deviations of the distribution of mean *R*_g_ values for each simulation temperature.

## Supporting Information

Please see *supporting information* for (a) the sequences shown by Garcia Quiroz and Chilkoti to have UCST and LCST phase behavior that were used for IS limit simulations in this study; (b) the temperature dependent free energies of solvation derived from free energy calculations using the AMOEBA forcefield; (c) figures shown the temperature dependent R_g_ profiles calculated in the IS limit; and (d) figures quantifying the statistics for residues selected as part of the design process directed toward the XPXXG system. A zip archive that is appended to the supporting information includes the parameter file for the AMOEBA forcefield and a sample key file for performing free energy calculations based on molecular dynamics simulations.

## Data Availability Statement

All of the data that support the findings in this study are available within the article and in the Supporting Information.

## Acknowledgments

This work was supported by grants DMR 1729783 from the US National Science Foundation (AC, RVP), RGP0034/2017 from the Human Frontier Science Program (RVP), and R01GM114237 from the US National Institutes of Health (PR). Resources from the Center for High Performance Computing (CHPC) and the Research Infrastructure Services (RIS) at Washington University in St. Louis were used for some of the simulations. XZ and RVP are grateful to Dr. Andreas Vitalis for several discussions over the years regarding issues that arise with regard to T-dependent free energies of solvation and the parameters for Δ*h* and Δ*c*_*P*_.

