## Supplemental Material for "Design of intrinsically disordered proteins that undergo phase transitions with lower critical solution temperatures"

**Table S1. Free energy of solvation of the model compounds at different temperatures calculated from AMOEBA simulations.** The statistical errors estimated from the BAR method, as described in detail by Wyczalkowski et al.,[1] are included in the brackets. The errors reported here are calculated as the root sum square of the statistical errors in each lambda window.

| Model Compound | $\Delta\mu_h(275K)$ kcal/mol | $\Delta\mu_h(298K)$ kcal/mol | $\Delta\mu_h(323K)$ kcal/mol | $\Delta\mu_h(358K)$ kcal/mol | $\Delta\mu_h(373K)$ kcal/mol | $\Delta\mu_h(398K)$ kcal/mol |
|---|---|---|---|---|---|---|
| methane | 1.25 (±0.04) | 1.63 (±0.04) | 1.98 (±0.04) | 2.06 (±0.03) | 2.29 (±0.03) | 2.29 (±0.03) |
| propane | 1.16 (±0.06) | 1.85 (±0.06) | 2.51 (±0.06) | 2.81 (±0.06) | 2.96 (±0.05) | 3.01 (±0.05) |
| 2-methylpropane | 1.46 (±0.07) | 2.22 (±0.07) | 2.87 (±0.07) | 3.11 (±0.07) | 3.24 (±0.06) | 3.35 (±0.06) |
| *n*-butane | 1.24 (±0.07) | 2.00 (±0.08) | 2.55 (±0.07) | 3.11 (±0.07) | 3.03 (±0.06) | 3.27 (±0.06) |
| ethyl methyl thioether | -2.60 (±0.08) | -1.92 (±0.08) | -1.24 (±0.08) | -0.89 (±0.07) | -0.46 (±0.07) | -0.26 (±0.06) |
| toluene | -0.97 (±0.09) | -0.17 (±0.09) | 0.46 (±0.09) | 0.75 (±0.08) | 1.11 (±0.07) | 1.14 (±0.06) |
| methanethiol | -1.45 (±0.05) | -1.04 (±0.05) | -0.72 (±0.05) | -0.42 (±0.05) | -0.15 (±0.05) | -0.11 (±0.04) |
| p-Cresol | -6.47 (±0.10) | -5.85 (±0.09) | -4.98 (±0.10) | -4.39 (±0.09) | -3.95 (±0.08) | -3.73 (±0.07) |
| 3-Methylindole | -5.49 (±0.12) | -4.46 (±0.12) | -4.04 (±0.11) | -3.77 (±0.10) | -3.30 (±0.09) | -3.26 (±0.08) |
| methanol | -5.40 (±0.05) | -5.08 (±0.05) | -4.59 (±0.05) | -4.18 (±0.04) | -3.79 (±0.04) | -3.50 (±0.04) |
| ethanol | -5.80 (±0.06) | -4.98 (±0.06) | -4.51 (±0.06) | -4.02 (±0.05) | -3.56 (±0.05) | -3.13 (±0.05) |
| acetamide | -8.94 (±0.07) | -8.61 (±0.06) | -8.03 (±0.06) | -7.59 (±0.06) | -7.22 (±0.06) | -6.79 (±0.05) |
| propionamide | -9.00 (±0.08) | -8.39 (±0.08) | -7.73 (±0.07) | -7.26 (±0.07) | -7.01 (±0.06) | -6.56 (±0.06) |
| 4-methylimidazole | -10.52 (±0.08) | -10.04 (±0.08) | -9.40 (±0.07) | -8.79 (±0.07) | -8.47 (±0.06) | -8.13 (±0.06) |
| N-methylacetamide | -9.09 (±0.07) | -8.33 (±0.08) | -7.81 (±0.07) | -7.24 (±0.07) | -6.83 (±0.06) | -6.34 (±0.06) |
| *n*-propylguanidine | -47.63 (±0.11) | -46.72 (±0.11) | -45.67 (±0.11) | -45.45 (±0.09) | -44.68 (±0.09) | -44.19 (±0.08) |
| 1-Butylamine | -61.22 (±0.10) | -60.49 (±0.09) | -59.53 (±0.09) | -58.99 (±0.08) | -58.28 (±0.08) | -57.63 (±0.07) |
| acetic acid (acetate) | -90.39 (±0.08) | -89.91 (±0.08) | -89.02 (±0.07) | -88.15 (±0.07) | -87.33 (±0.07) | -86.32 (±0.06) |
| propionic acid (propionate) | -86.84 (±0.12) | -86.16 (±0.12) | -85.24 (±0.12) | -84.32 (±0.11) | -83.35 (±0.10) | -82.61 (±0.07) |

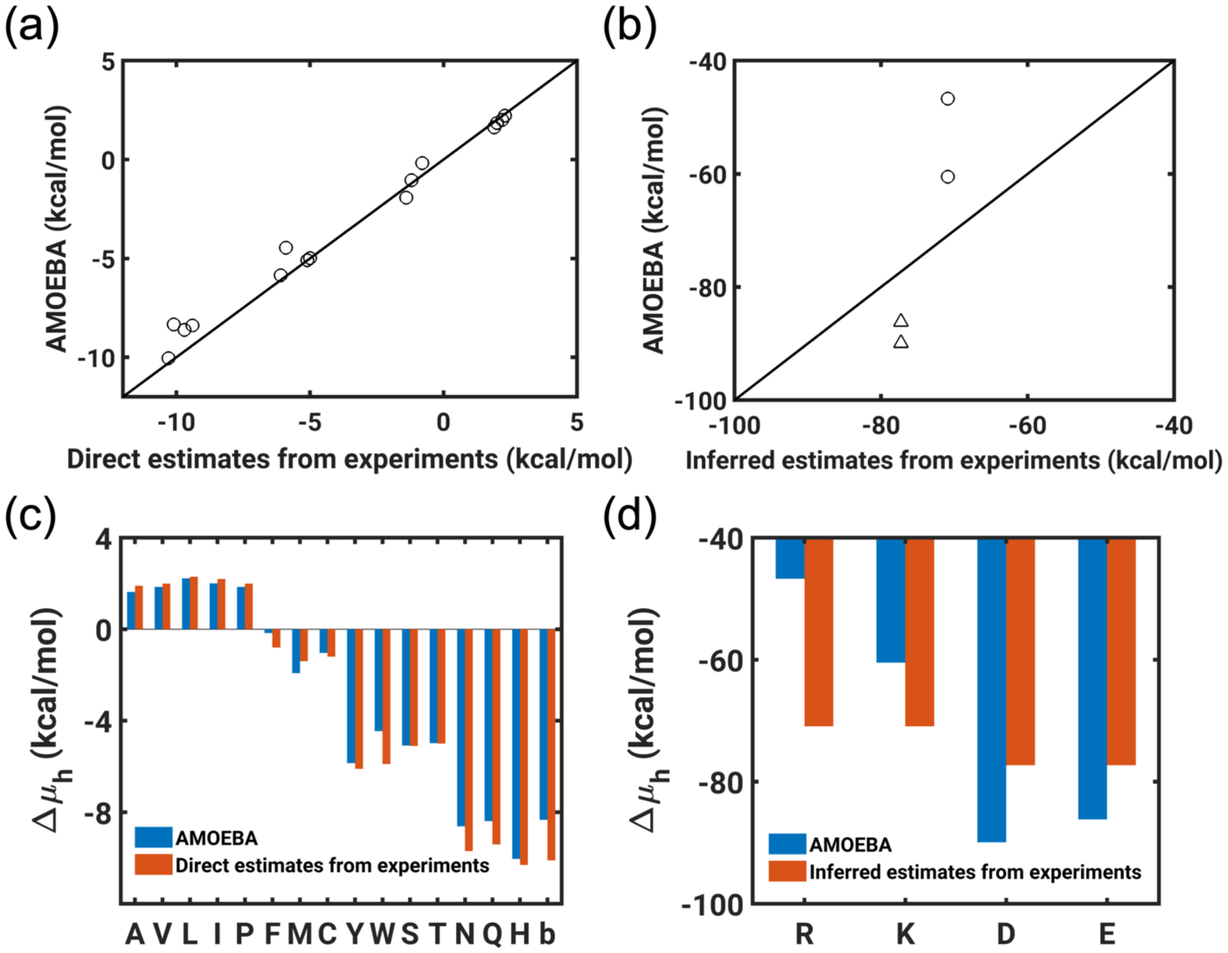


**Figure S1. Comparison between the free energy of solvation calculated from AMOEBA simulations at 298 K and experimentally derived values used to date in ABSINTH [2].** (a) Comparisons for neutral model compounds; (b) Comparisons for charged model compounds; the acidic groups and basic groups are shown as triangles and circles, respectively. Bar plots of these data are also shown for (c) neutral model compounds and (d) charged model compounds. Here, 'b' in panel (c) refers to the backbone moiety, modeled using N-methylacetamide (NMA), that mimics the peptide unit. It is worth noting that the oft-quoted value for NMA is -10 kcal / mol [3]. These values have been revisited a few times and the Guthrie hydration free energy database (https://escholarship.org/uc/item/53n2h10t) [4] cites eight separate measurements, and lists values of -10.1 kcal / mol, -10.0 kcal / mol, -9.9 kcal / mol, -9.0 kcal / mol, -7.3 kcal / mol, with an overall error of 1.93 kcal / mol. Results in ABSINTH-based simulations are typically insensitive to changes by ± 2 kcal / mol for polar moieties[5]. Whether the discrepancy for Trp is significant enough to warrant corrections remains unresolved at this juncture. We emphasize that the primary focus behind using AMOEBA-based numbers in our work was to obtain direct estimates for charged moieties. We use AMOEBA-based numbers for all other groups to maintain consistency across functional groups.

**Table S2. Sequences that show LCST and UCST phase behavior. From the work of Garcia Quiroz and Chilkoti [6].** Each sequence is preceded by a numeric ID.

| LCST | | | UCST | |
|---|---|---|---|---|
| 1: $(APVGVG)_{10}$ | 8: $(TVPGVG)_{9}$ | 15:$(VTPAVG)_{9}$ | 20: $(GRGNSPYG)_{7}$ | 26: $(GRGDNPYQ)_{6}$ |
| 2: $(VAPVG)_{11}$ | 9: $(GVPGAV)_{9}$ | 16:$(VHPGVG)_{9}$ | 21: $(GRGDSPYG)_{7}$ | 27: $(QYPSDGRG)_{6}$ |
| 3: $(VPAVG)_{11}$ | 10: $(GVGGVA)_{9}$ | 17:$(VPHVG)_{11}$ | 22: $(GRGDSPFG)_{7}$ | 28: $(RGDSPYG)_{7}$ |
| 4: $(TPVAVG)_{8}$ | 11: $(VPGVA)_{11}$ | 18:$(VGPAVG)_{9}$ | 23: $(GRGESPYG)_{7}$ | 29: $(RGDSYPG)_{7}$ |
| 5: $(VPSALYGVG)_{5}$ | 12: $(VGPVG)_{11}$ | 19:$(VPSTDYGVG)_{6}$ | 24: $(RGDSPYQG)_{7}$ | 30: $(RGDSPHG)_{8}$ |
| 6: $(VPGVG)_{11}$ | 13: $(VPAGVG)_{9}$ | | 25: $(GRPSDSYG)_{7}$ | |
| 7: $(AVPGVG)_{9}$ | 14: $(VPTGVG)_{9}$ | | | |

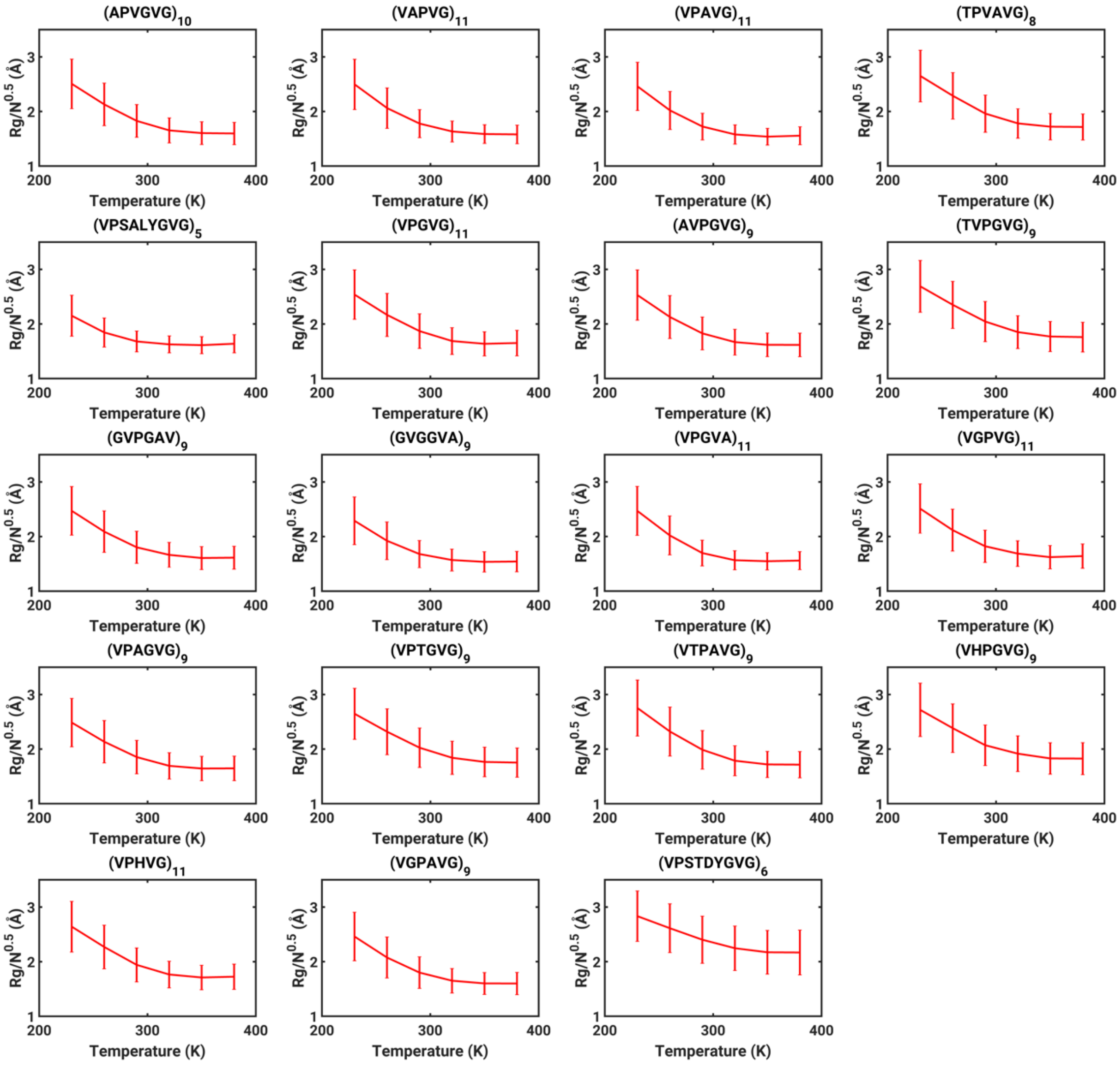


**Figure S2. Normalized $R_g$ values as a function of simulation temperatures for the sequences in Table S2.** The simulations are based on the IS limit of the ABSINTH model. The error bar indicates the standard derivation of the $R_g$ distribution. In general, the IS limit simulations were highly reproducible, and therefore, for each construct, we performed one thermal replica exchange simulation comprising $6\times10^7$ MC steps per replica. The temperature schedule ranges uniformly from 230 K to 380 K in steps of 30 K.

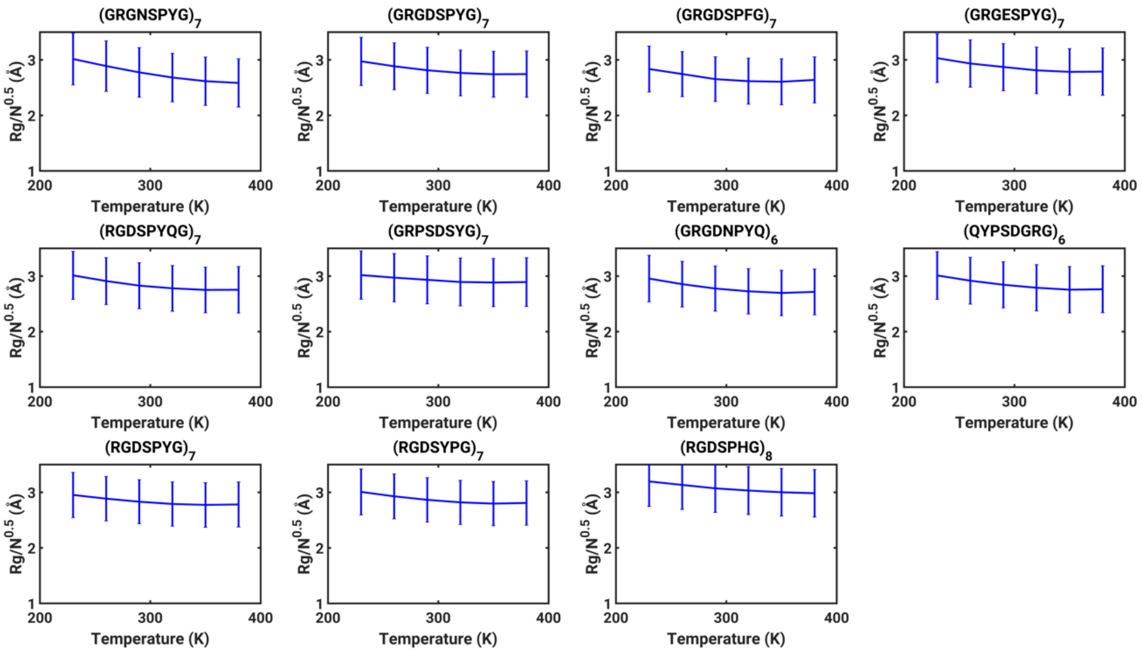


**Figure S3. Normalized $R_g$ values, derived from IS limit simulations, plotted against simulation temperatures for the UCST sequences in Table S2**. As with **Figure S2**, the error bars indicate standard derivations of the $R_g$ distribution. The details of the thermal replica exchange simulations are as described for the sequences that show LCST behavior. Notice the near zero slopes for $R_g$ versus T in contrast to the situation for the LCST sequences shown in **Figure S2**.

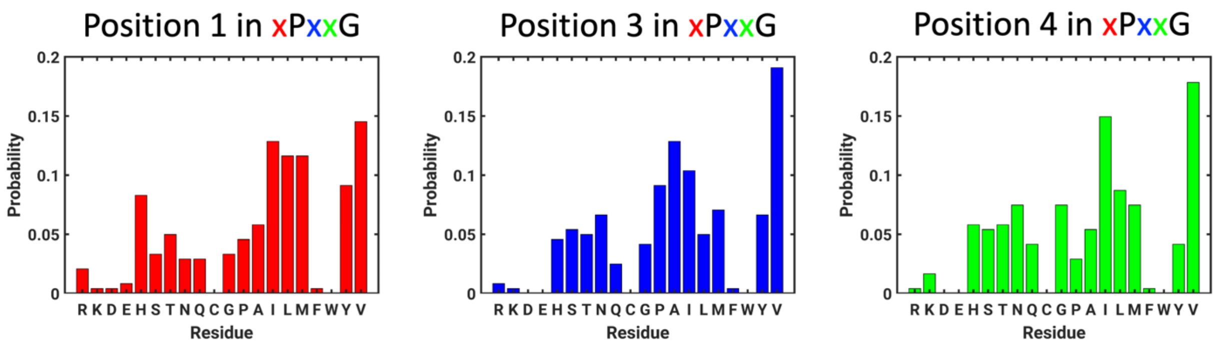


**Figure S4. Position-specific probabilities of finding each of the nineteen non-Cys residues at the three positions, 1, 3, and 4 that are mutable in an xPxxG sequence for each repeat.** It takes six iterations of the genetic algorithm to generate 241 distinct sequences with $m < 5\times10^{-3}$ åK$^{-1}$.
